# The effects of iron limitation on the small chlorophyte *Micromonas* from the Northeast Pacific Ocean

**DOI:** 10.1101/2024.09.26.615135

**Authors:** Meredith G. Meyer, Vincent J. White, Olivia Torano, Heidi Hannoush, Margarita Lankford, Adrian Marchetti

## Abstract

Small eukaryotic phytoplankton can account for a considerable amount of biomass and primary production in high nutrient, low chlorophyll (HNLC) regions of the ocean where iron-limitation is pronounced. However, the physiological and metabolic strategies these cells invoke to cope under low iron conditions and the extent to which they are responsible for new production (i.e., the fraction of primary production supported by nutrients from outside of the euphotic zone) is unclear. Here, we examine how a representative picoeukaryote, the chlorophyte *Micromonas* sp., recently isolated from the iron-limited subarctic Northeast Pacific Ocean, responds to iron-limitation when grown on nitrate as a nitrogen source. Iron-limited *Micromonas* exhibits reductions in growth rate, cell volume, and elemental quotas along with a restructuring of cellular metabolism. Gene expression and pathway analyses show evidence of strategies to mitigate iron limitation with constitutive expression of genes related to nitrogen uptake and utilization. Additionally, cellular carbon and nitrogen quotas ranged from 20 – 170 fmol C cell^−1^ and 3.3 – 20 fmol N cell^−1^, respectively, as a function of iron status. Based on the measured cellular quotas, we estimate that representative picoeukaryotes (<2 µm), such as *Micromonas,* in HNLC Northeast Pacific waters can account for a significant proportion of new production, supporting the need for a reconsideration of the role small eukaryotic phytoplankton play in the global carbon cycle.

## 1. Introduction

In open-ocean ecosystems, high surface area-to-volume ratios and generally lower nutrient quotas make picophytoplankton strong competitors for nutrients, thus enabling them to routinely dominate eukaryotic phytoplankton communities in these oligotrophic environments (Worden and Not, 2008). While much research has focused on cyanobacteria, specifically *Prochlorococcus* (Chisholm et al., 1991) and *Synechococcus* (Moore et al., 2002), fewer studies have investigated the growth strategies of picoeukaryotes which have been shown to potentially play key ecological roles (Fowler et al., 2020). Previous studies have suggested chlorophytes and other small eukaryotic groups primarily rely on ammonium and other forms of recycled nitrogen to support their growth (Greene et al., 1992; Marchetti et al, 2006; Meyer et al., 2022). Compared to cyanobacteria, diatoms, and dinoflagellates, these picoeukaryotes are poorly characterized and are less likely to be available in culture. Thus, there are fewer monocultured representatives sequenced for genomic and/or transcriptomic analyses compared to other phytoplankton groups (Le Gall et al., 2008). This overall lack of knowledge has led to these organisms commonly being overlooked in whole community analyses despite their high abundances.

Given the potentially outsized role these eukaryotic picophytoplankton could play in oligotrophic and nutrient-limited regions, understanding the nutrient acquisition strategies, requirements and metabolisms employed by these phytoplankton and the resultant effect for general ecosystem productivity and carbon export potential is of substantial importance. Here we present measurements obtained from the chlorophyte, *Micromonas* sp., that was recently isolated from Ocean Station Papa (Station P) in the subarctic Northeast Pacific during the NASA EXport Processes in the Ocean from RemoTe Sensing (EXPORTS) program in 2018. *Micromonas* sp. is an abundant member of the *Mamiellales* order and is closely related to *Ostreococcus* and *Bathycoccus* (Monier et al., 2016; Vannier et al., 2016), two other ecologically important picoeukaryotic chlorophytes. Studies have shown that *Micromonas* can occupy a wide range of habitats, from the tropics to the poles as well as offshore and coastal environments (Demory et al., 2018). The group often exhibits an “occupy-abundance pattern” where *Micromonas*, and other members of the *Mamiellophyceae* lineage, frequently become the most abundant genera in regions they are present (Monier et al., 2016). Yet, the metabolic capabilities and the ecological role of *Micromonas* in iron-limited regions is still poorly understood.

Station P is the site of a long-term oceanographic monitoring program which resides within a High Nutrient (predominantly nitrate (NO_3_^−^)), Low Chlorophyll, or HNLC, region. This region is chronically limited by the bioavailability of iron which, despite replete macronutrient concentrations, inhibits substantial phytoplankton biomass standing stock and higher rates of primary production (Martin et al., 1988; Marchetti et al., 2006). Iron is used as a crucial co-factor in photosynthesis and nitrogen assimilation (LaRoche et al.,1993; Peers and Price, 2006). Many groups of phytoplankton that experience iron limitation have been shown to utilize regenerated nitrogen sources (e.g., ammonium (NH_4_^+^) or urea) which theoretically have a lower iron requirement relative to oxidized forms of nitrogen or nitrogen gas (Price et al., 1994; Varela and Harrison, 1999). In addition, recent findings suggest that the most dominant ecotype of the cyanobacteria in the Northeast Pacific, *Synechococcus*, lacks the ability to assimilate NO_3_^−^, thus being restricted to regenerated production (i.e., using recycled nitrogen sources such as ammonium and urea) (Sharpe et al., 2023). This finding presents a knowledge gap in our understanding of which organisms are responsible for the vast majority of new production in the region, which has routinely been measured to be driven predominantly by small phytoplankton groups (i.e., < 5 μm in cell diameter) (Meyer et al. 2022).

Here, we sought to examine why *Micromonas* can be found in high abundances in the Northeast Pacific and their possible role in carbon export. We had two primary research objectives: a) examine the distinct physiological and metabolic strategies of *Micromonas* to grow under low iron conditions and b) quantify the extent to which *Micromonas* can contribute substantially to new production in this region. We performed these objectives through determining rates of cellular growth, photophysiology, carbon and nitrogen cellular content, and trends in gene expression as a function of iron status. The combination of molecular and physiological analyses provides a more wholistic approach to examining ecologically important phytoplankton groups and their influence on marine biogeochemical cycles.

## 2. Methods

### 2.1 Sampling and Cell Isolation

Whole seawater was collected from approximately 5 m depth via a trace metal clean rosette during the NASA EXPORTS field campaign to Ocean Station Papa (Station P) in the subarctic North Pacific. This campaign occurred from August 16^th^ to September 7^th^, 2018. To keep storage conditions as similar to the ambient environment as possible, sampled seawater was placed in culture flasks and incubated in on-deck incubators before being transported back to the University of North Carolina for phytoplankton isolation. An abundant, small (< 5 µm) chlorophyte was isolated via single-cell isolation using a 10 μL pipette. DNA was extracted using standard protocols with a DNEasy Plant Mini Kit (Qiagen). The V4 region of the 18S rRNA gene was sequenced (National Center for Biotechnology Information (NCBI) ID PRJNA1072555), and the consensus sequence of forward and reverse reads was searched against the NCBI repository using BLAST, revealing the highest sequence similarity to *Micromonas pusilla* (accession number KY980282.1) with a 100% percent ID and 76% query cover.

### 2.2 Culture conditions

*Micromonas sp*. (UNC1837) cells were maintained in Aquil media in an incubator under constant 120-150 µmol photons m^−2^ s^−1^ light at 12°C to maintain light saturation and ambient temperatures close to environmental conditions at Station P. To examine the effects of differing nitrogen forms under variable iron states on *Micromonas* growth and physiology, four trace metal clean culture treatments were initiated. Aquil medium was sterilized in a microwave, cooled, and supplemented with filter-sterilized (0.2-μm Acrodisc) ethylenediamine tetraacetic acid (EDTA) trace metals (minus iron), vitamins (B_12_, thiamine, and biotin), and chelexed macronutrients (NO_3_^−^, phosphate) in accordance with Aquil medium concentrations (Price et al. 1989). Trace-metal concentrations were buffered using 100 µM of EDTA. Two treatments contained NO_3_^−^ (300 µM) and two treatments contained NH_4_^+^ (30 µM) as their nitrogen source. Within each nitrogen treatment, there was an iron replete (1370 nM total Fe) and an iron deplete (3.1 nM total Fe) treatment. Iron concentrations were selected after initial testing, and the chosen concentrations were able to achieve the most consistent growth rates after a minimum of three transfers with transfers occurring approximately every 7-10 days. Thus, the four treatments can be described as NO_3_^−^ Fe+, NO_3_^−^ Fe−, NH_4_^+^ Fe+, and NH_4_^+^ Fe−. Each treatment was first cultured in triplicate acid-washed 28 mL clear, polycarbonate centrifuge tubes (Nalgene, Rochester, NY) for acclimation to their respective treatments. Cells were acclimated to their treatment for a minimum of three transfers before experiments began. During acclimation, 200 µL of dense late exponential-phase culture was transferred to new tubes with fresh media to maintain growth, thus representing a 125-fold dilution. Following acclimation, cultures were transferred to a trace metal clean, 1 L polycarbonate bottle that was approximately 700 mL full. We were not able to maintain steady state growth rates in NH_4_^+^ Fe+ and NH_4_^+^ Fe− cultures, thus confining our experiment to the NO_3_^−^ Fe+ and NO_3_^−^ Fe−, referred to as Fe+ and Fe− treatments herein. All culture preparation, maintenance, and cell transfers were conducted in a trace metal clean lab under a positive-pressure, laminar flow hood.

During the experiments, approximate cell growth and concentrations were monitored via relative fluorescence units (RFUs) on a Turner 10AU fluorometer from lag phase through stationary phase (Brand et al. 1981). Once samples were determined to be in mid-exponential phase, cultures were measured for RFUs and other photophysiological parameters using a Satlantic FIRe fluorometer with blue light diode (450 nm) and 60 iterations per triplicate measurement (see below) and harvested for the following parameters: cell counts, chlorophyll *a* (Chl *a*) concentration, particulate carbon (C) and nitrogen (N) concentration, and RNA.

Specific growth rates (µ) were calculated as the slope of the natural log of relative fluorescence units over time, according to Brand et al. (1981). Assuming Fe+ represents a maximum growth rate (µ_max_), the ratio of average µ/µ_max_ (i.e., Fe−/Fe+) can be calculated. The maximum quantum yield of photochemistry in photosystem II (PSII; F_v_/F_m_) and the functional absorption cross section of PSII (σ_PSII_) were measured as metrics to evaluate photosynthetic efficiency (Gorbunov and Falkowski; 2004).

Chl *a* was analyzed fluorometrically following Graff and Rynearson (2011). Particulate carbon (PC) and nitrogen (PN) were analyzed on a Carlo Erba NC 2500 elemental analyzer at University of Maryland Center for Environmental Science Appalachian Laboratory in Frostburg, Maryland. Cultures for RNA were filtered onto an Isopore 0.4 µm pore size, 47 mm polycarbonate filter under gentle vacuum pressure, flash frozen in liquid nitrogen, and stored at −80 °C before extraction. RNA was extracted using standard protocols with a RNAqueous-4PCR kit according to the manufacturer’s protocols. Extracted RNA was checked for quality (A280/260, A260/230, and nucleic acid concentration) on a Nanodrop spectrophotometer and cleaned using a RNeasy MinElute Clean-up kit. RNA was sent to the Azenta Genewiz Sequencing Facility in South Plainfield, New Jersey for Illumina high throughput analysis with ∼350M raw paired-end reads per lane and single index, 2 x 150 bp per lane, resulting in a read depth of 62-65 million reads per sample.

### 2.3 Cell counts and size estimates

A small aliquot (1.8 mL) of each culture (both treatments and all replicates) were preserved in 200 µL of paraformaldehyde in 2 mL cryovials at the time of cell harvest, the day following the harvest and several (2-3) days preceding the harvest. Cryovials were immediately flash frozen in liquid nitrogen and stored at −80 °C until they were analyzed on a Guava easyCyte flow cytometer equipped with a red laser in the Paerl Lab at North Carolina State University. Data were processed by InCyte software to provide cell counts based on red fluorescence. Estimates of cell size (diameter) were also calculated using calibration beads of known size based on the relationships of Paerl et al. (2020). Cell volume was calculated by assuming cells were spherical. Cellular Chl *a*, PC and PN concentrations were calculated by normalizing concentrations by cell counts and by cell biovolume.

### 2.4 North Pacific new production calculations

If we assume *Micromonas* is a good representative (i.e., in the top 20% most represented species in 18S rRNA gene community composition data; Jones and Rynearson, 2022) of the picoeukaryote population at Station P, upper bound estimates of the contribution of picoeukaryotes to total new production under Fe− replete and Fe− limiting conditions can be calculated according to Equation 2:

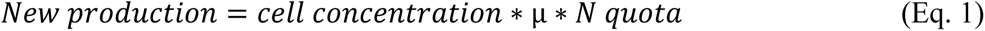

Where the nitrogen quota is in units of fmol N cell^−1^. Here, the N quota and growth rate (µ; d^−1^) come from our laboratory results with the *Micromonas* culture. The cell concentration used in Fe+ and Fe− culture new production rates is the picoeukaryote cell counts (3.3 × 10^6^ cells L^−1^) measured from the EXPORTS Northeast Pacific cruise in 2018 (Graff, 2023). These are average picoeukaryote counts collected from surface waters and measured via flow cytometry during the observation period.

### 2.5 Gene expression analysis

Transcripts were trimmed of sequence adapters and poly-A tails via Trim_galore (v.0.6.2; Altos Labs) and underwent quality control analysis via FastQC (v.0.11.9) where sequences with a quality score below 20 and/or length less than 50 base pairs were removed. Merged sequence files were assembled de novo via Trinity (v.2.8.6) before being combined into grand assemblies based on iron treatment via cdhit (v.4.8.1; Fu et al., 2012). Sequences underwent functional annotation via the Kyoto Encyclopedia of Genes and Genomes (KEGG; Release 88.2). Additional manual annotation of genes associated with Fe metabolism, but are not included in KEGG (e.g., protin-pumping rhodopsins, iron starvation induced proteins (ISIPs), and phytotransferrins), was conducted (see Moreno et al., 2017 for more information on manual annotation and sequence acquisition). Independent end-to-end read alignment occurred via Salmon (v.10.9.1; Patro et al., 2017) and annotation and count files were combined via tximport, providing counts per million (CPM) values per gene per sample (Soneson et al., 2015). Differential expression of genes was analyzed at the KEGG Ortholog (KO) level, averaging values for multiple contigs with the same KO, according to the DESeq2 package in RStudio (v.4.2.2). The Log2Fold change (adjusted p-value < 0.05) of specific genes of interest, particularly those associated with photosynthesis, N metabolism, and Fe metabolism, were investigated. Additionally, trends in KEGG metabolic pathways were analyzed among samples. Figures were generated in Matlab R2022b. Sequences from this analysis are available at the National Center for Biotechnology Information (Ascension numbers: SAMN39858273 - 284).

## 3. Results

### 3.1 Growth rates and photosynthetic efficiency

The Fe+ cultures had an average growth rate of 0.96 ± 0.01 d^−1^ (Fig. 1A). The average growth rate of Fe− cultures was substantially lower at 0.48 ± 0.07 d^−1^ resulting in a µ/µ_max_ (Fe−/Fe+) of 0.50, suggesting substantial limitation of iron in the Fe− treatment relative to the Fe+ treatment (Fig. 1A). Estimates of maximum photochemical yield of PSII (F_v_/F_m_) confirm that the Fe+ cultures had an increased efficiency of PSII compared to Fe−, which was experiencing iron-limited conditions. The mean F_v_/F_m_ for the Fe+ treatment was 0.73 ± 0.02, whereas values for the Fe− cultures were 0.37 ± 0.03, consistent with a decline in photosynthetic efficiency of PSII due to iron limitation (Fig. 1B). The cross-sectional absorption area of photosystem II (σ_PSII_) was also consistent with other iron-limited phytoplankton cultures with an increase in σ_PSII_ under iron-limited conditions (Ellis, 2015; Moreno et al., 2020). For Fe− cultures, σ_PSII_ averaged 1036 ± 264, and for Fe+, σ_PSII_ averaged 372 ± 12 (data not shown), resulting in a 2.8-fold increase under iron-limited conditions.

**Fig. 1.**
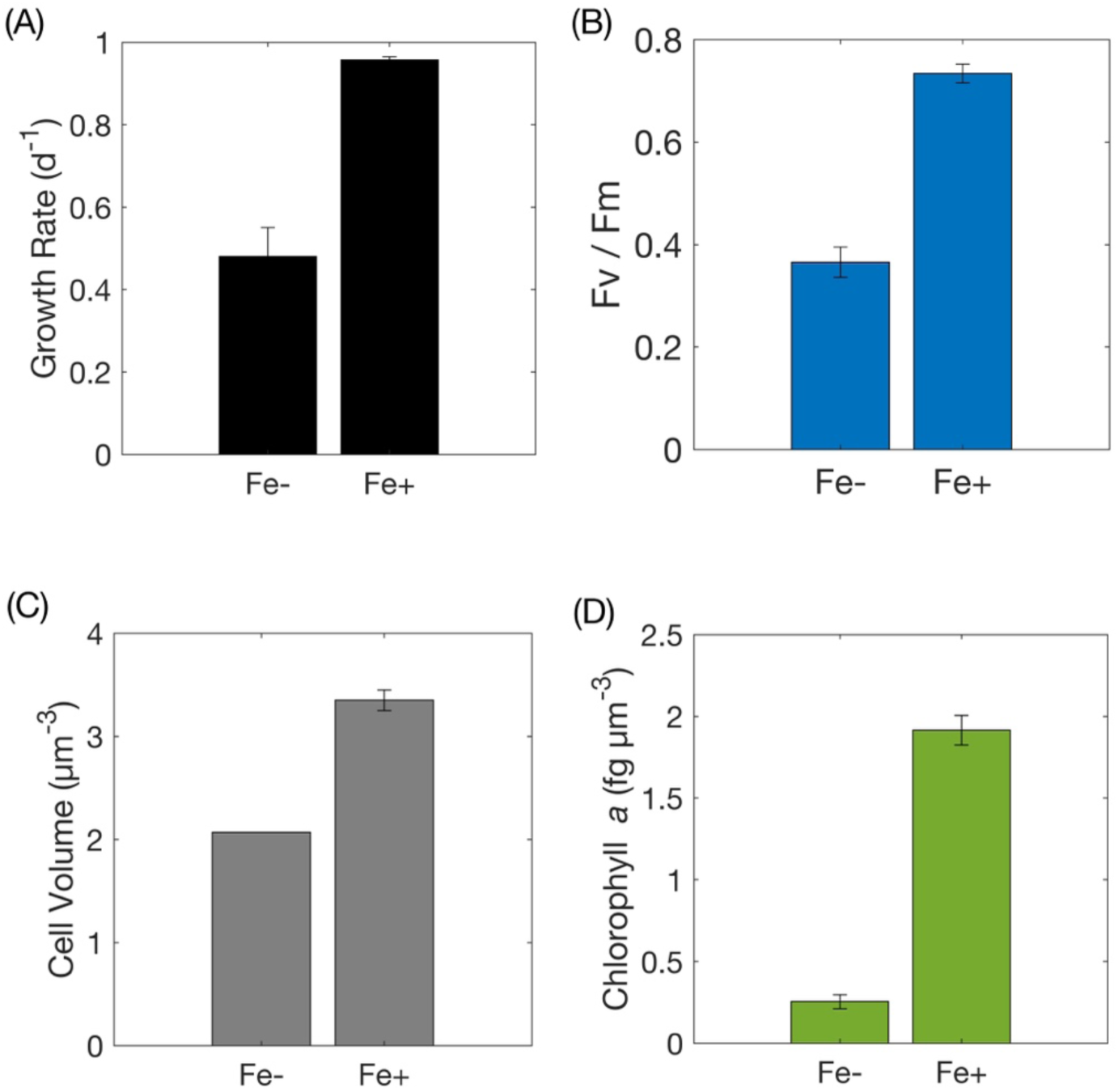
Specific growth rates (A; d^−1^), maximum photochemical efficiencies of PSII (B; F_v_/F_m_), cell volumes (C; µm^3^), and cellular chlorophyll *a* quotas (D; fg µm^−3^) for *Micromonas* grown under iron-replete (Fe+) and iron-limited (Fe−) conditions. Error bars represent standard deviations with n = 3 for Fe− and n = 2 for Fe+.

There was a substantially lower F_v_/F_m_ value for the Fe+ replicate A sample compared to the other replicates, suggesting that this culture may have entered stationary phase, experiencing N depletion at the time of sampling. This triplicate was therefore removed from the averages of Fe+ sample data presented in Figures 1 and 2 as well as from the gene expression analysis. Although with all replicates included, there was still a significant difference in growth rates, cellular Chl *a* concentrations and cellular C and N quotas between the Fe+ and Fe− treatments (Supplemental Table 1).

**Fig. 2.**
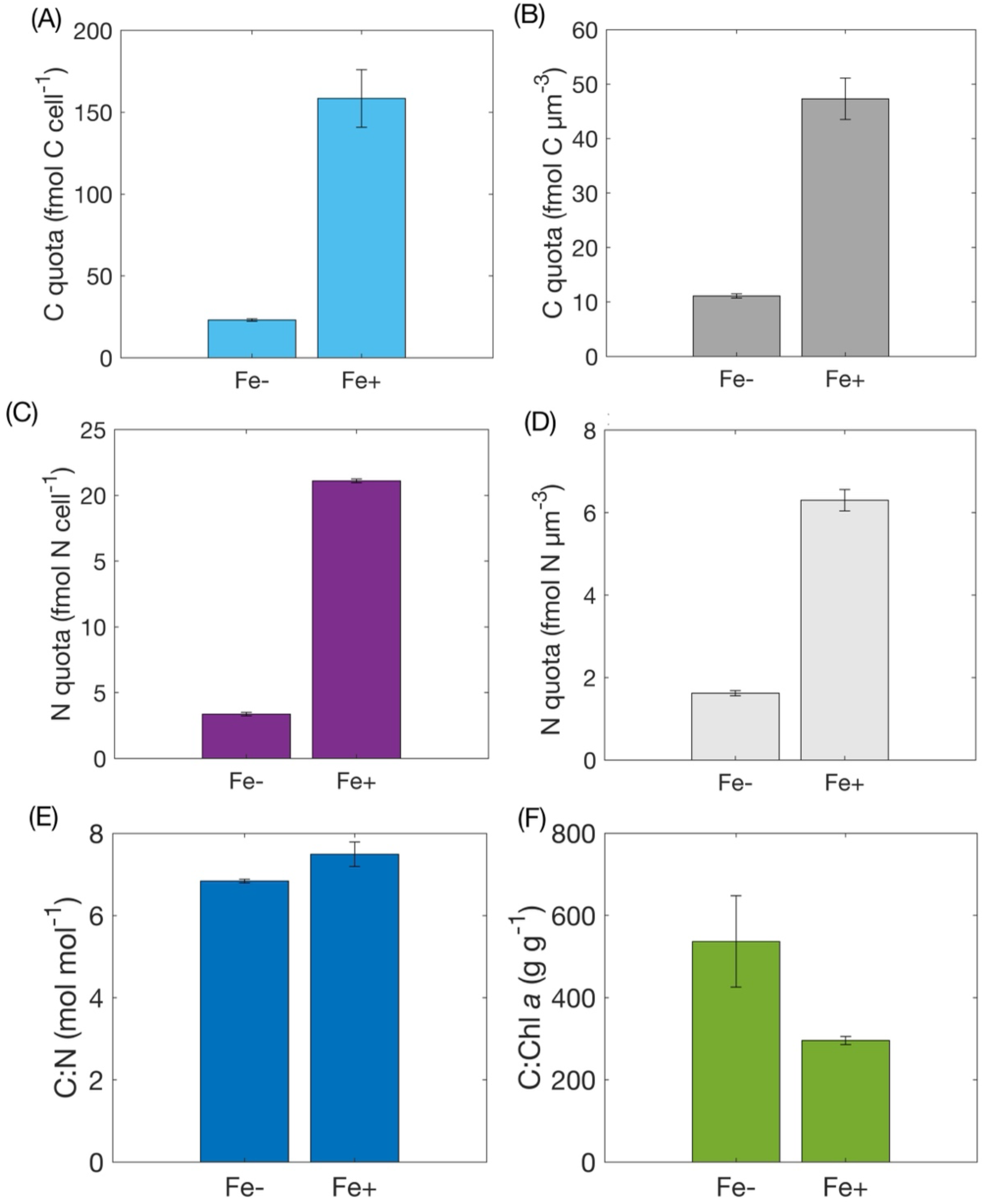
Cellular carbon (A,B) and nitrogen (C,D) quotas and C:N (E) and C:Chl *a* (F) ratios for *Micromonas* grown under iron-replete (Fe+) and iron-limited (Fe−) conditions. Quotas are represented per cell (A,C; fmol cell^−1^) and per cell volume (B,D; fmol µm^−3^). Error bars represent standard deviations with n = 3 Fe− and n = 2 for Fe+.

### 3.2 Cell Size and chlorophyll a content

The Fe+ cultures had an average cell diameter of 1.85 ± 0.02 µm (data not shown) and corresponding cell volume of 3.35 ± 0.10 µm^3^ (Fig. 1C). The Fe− cultures had an average cell diameter of 1.58 ± 0.00 µm and cell volume of 2.07 ± 0.00 µm^3^ (Fig. 1C).

Cellular Chl *a* contents exhibited an order of magnitude difference between the Fe+ and Fe− samples, with the Fe+ cultures having an average of 6.42 ± 0.50 fg cell^−1^ and the Fe− cultures averaging only 0.53 ± 0.09 fg cell^−1^. Chl *a* concentration, when normalized to cell biovolume, exhibited a similar trend to cellular Chl *a* content, but the difference in the magnitude between the Fe+ and Fe− cultures was substantially lower (Fig. 1D). The average biovolume-normalized Chl *a* content was 1.92 ± 0.09 fg µm^−3^ in the Fe+ treatment and 7.5-fold lower at 0.25 ± 0.04 fg µm^−3^ in the Fe− treatment.

### 3.4 Cellular carbon and nitrogen quotas

Like Chl *a*, the cellular C content exhibited an order of magnitude difference between the Fe+ and Fe− treatments. The Fe+ cultures had a high cellular C quota of 158 ± 20 fmol C cell^−1^ whereas the Fe− cultures had an 8-fold lower C quota of 23 ± 0.0 fmol C cell^−1^ (Fig. 2A). As is seen throughout the parameters measured, the Fe− sample quotas were consistent, only ranging from 22 – 24 fmol C cell^−1^ among triplicates. When normalized to cell volume, the biovolume-normalized C quotas were more similar between treatments due to the cells in the Fe+ culture being ∼1.5-fold larger than the cells in the Fe− culture. The Fe+ treatment had a C quota of 47 ± 3.8 fmol C µm^−3^ whereas the quota for the Fe− treatment was 11.1 ± 0.38 fmol C µm^−3^ (Fig. 2B). Thus, when considering the difference in cell volumes (i.e., calculated per µm^−3^ rather than per cell), the difference in C quota decreases from 8.0- to 4.3-fold higher in the Fe+ treatment relative to the Fe− treatment.

The cellular N contents likewise exhibited orders of magnitude differences between iron treatments when normalized per cell and per unit volume. The average N quota for the Fe+ cultures was 21.1 ± 1.5 fmol N cell^−1^ whereas the average for the Fe− cultures was 3.37 ± 0.13 fmol N cell^−1^ (Fig. 2C). The per unit volume average for Fe+ samples was 6.30 ± 0.26 fmol N µm^−3^, and for the Fe− samples was 1.62 ± 6.44 × 10^−2^ fmol N µm^−3^ (Fig. 2D). The change in cellular N contents as a function of iron status was similar to that of C at approximately 6- and 4-fold larger for the Fe+ treatment when normalized per cell and per unit volume, respectively. Both treatments exhibited slightly higher particulate carbon : nitrogen (C:N) ratios than the Redfield ratio of 6.6 with averages of 7.49 ± 0.30 and 6.84 ± 0.05 for Fe+ and Fe−, respectively (Fig. 2E). C:Chl *a* ratios exhibited the opposite pattern with a higher average ratio (537 ± 111 µg : µg) in the Fe− treatment than in the Fe+ treatment (296 ± 9.74 µg : µg; Fig. 2F).

### 3.5 Potential contributions to new production in the NE Pacific

The average picoeukaryote cell concentration during the EXPORTS observation period used in these calculations was 3.34 × 10^6^ cells L^−1^ (Graff, 2023). Based on these picoeukaryote cell densities, rates of potential contributions to new production varied by an order of magnitude between iron treatments, averaging 5.4 ± 1.0 nmol N L^−1^ d^−1^ for Fe− cultures and 67.5 ± 4.3 nmol N L^−1^ d^−1^ for Fe+ cultures (Fig. 3). Rates were highly variable, with Fe− values ranging from 3.7 to 8.4 nmol N L^−1^ d^−1^ and Fe+ values ranging from 45.9 to 105.1 nmol N L^−1^ d^−1^. Rates of small-celled new production at Station P were always between the Fe− and Fe+ rates, except for on YD 240 when the EXPORTS rate was higher than Fe+ (55.4 nmol N L^−1^ d^−1^) at 58.5 nmol N L^−1^ d^−1^.

**Fig. 3.**
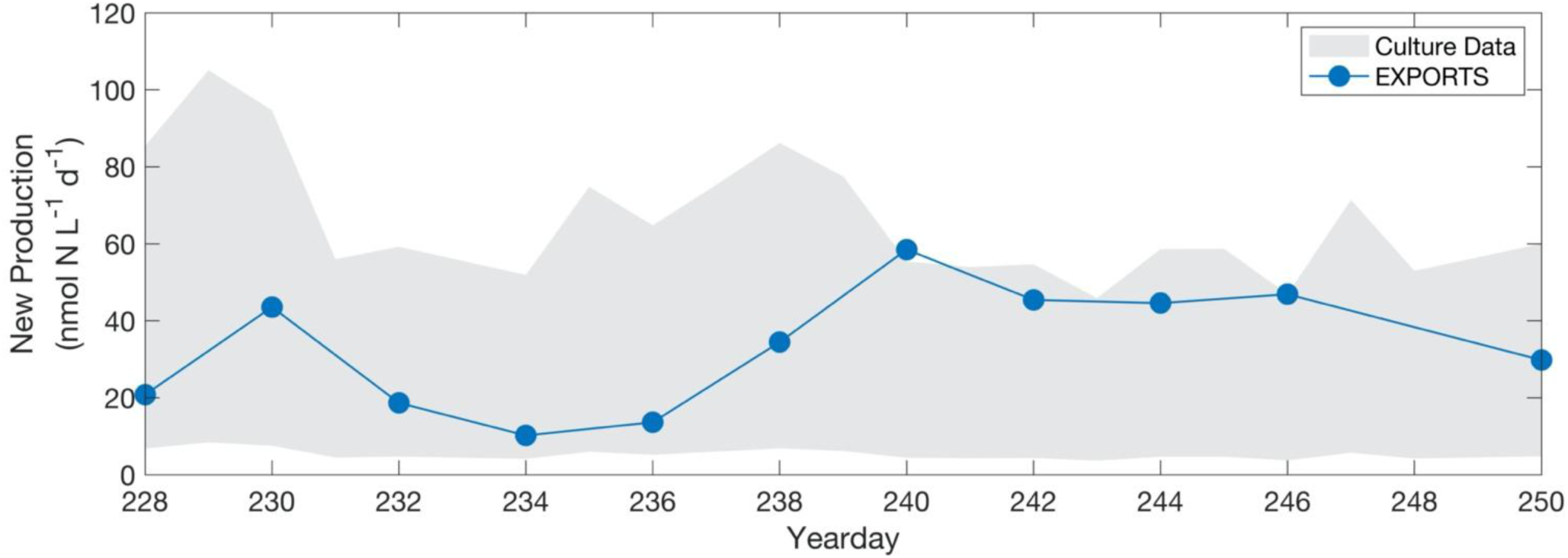
Estimates of *Micromonas sp.* contributions to new production during the North Pacific EXPORTS cruise. *Micromonas sp.* new production rate estimates for NO_3_^−^ Fe− and NO_3_^−^ Fe+, are calculated from culture-based growth rates, cellular N quotas, and in situ NP EXPORTS picoeukaryotic cell counts obtained from flow cytometry. The gray shaded region represents the upper bounds (NO_3_^−^ Fe+) and lower bounds (NO_3_^−^ Fe−) for the culture-based rate estimates. The blue line represents the average mixed-layer rates of new production of the small the size-fraction (< 5 µm) during EXPORTS by yearday during the observation period.

### 3.6 Gene expression

Molecular sequencing of the *Micromonas* sp. (UNC1837) transcriptome indicated a higher GC content for the Fe+ (61%) vs. Fe− (49%) cultures. Total sequences and sequence length were comparable between treatments for the Fe+ and Fe−, respectively.

The impact of Fe availability was analyzed broadly on the metabolic pathway level (Fig. 4A) and specifically on a targeted gene level (Fig. 4B). Of the 17 metabolic pathways of interest analyzed, on average, 10 displayed overrepresentation in Fe− treatment (i.e., negative Log2Fold change). Key pathways exhibiting this trend include those related to iron limitation mitigation strategies or general physiological stress responses, including iron starvation induced protein 3 (ISIP3) expression, ferredoxin expression, higher expression of photosystem I relative to photosystem II, and zeaxanthin epoxidase (Fig. 4A). Unsurprisingly, many gene transcripts involved in pathways related to general cell maintenance, including nitrogen metabolism, arginase, flavin adenine dinucleotide synthesis, etc., were overrepresented in the Fe+ treatment.

**Fig. 4.**
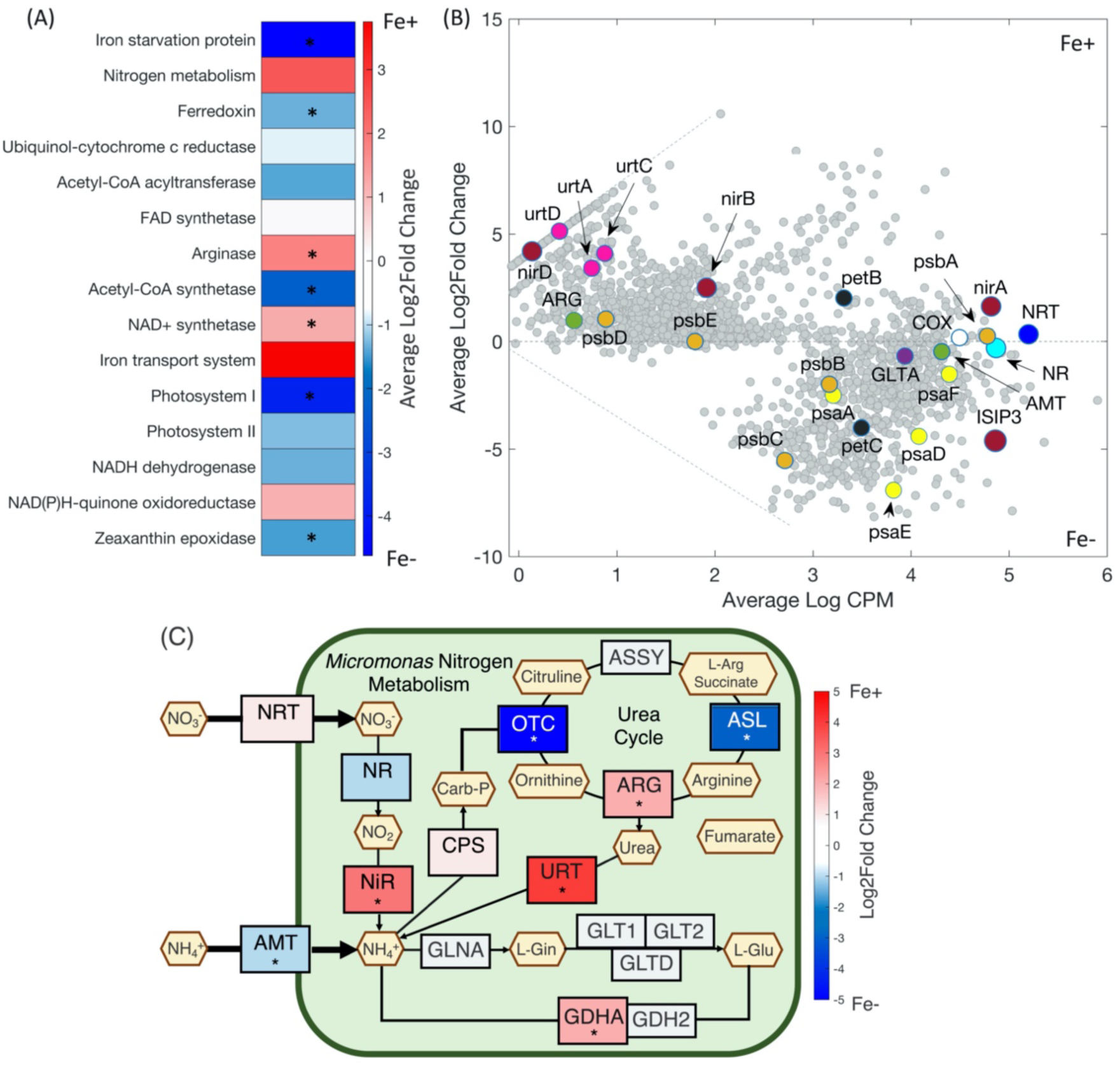
*Micromonas sp.* gene expression as a function of iron status. Heatmap (A) of average Log2Fold change of KEGG pathways, RA plot (B) of average log counts per million (CPM) vs. average Log2Fold change, and nitrogen assimilation pathway map (C) of *Micromonas sp.* Fe+ vs. Fe− culture RNA transcripts. Transcripts were annotated and evaluated on the KO level using a significance threshold of p <0.05. For all plots, positive Log2Fold change indicates overrepresentation in Fe+ cultures and negative Log2Fold change indicates overrepresentation in Fe− samples. Astericks on the heatmap and pathway map indicate significant changes (p <0.05). The RA highlights protein-encoding genes of interest. The cellular pathway map indicates the sign and relative strength of Log2Fold change expression of genes. Grey genes are absent from transcript data. Abbreviations are: nitrate transporter (*NRT*), nitrite reductase (*NiR*), nitrate reductase (*NR*), ammonium transporter (*AMT*), ornithine carbamolytransferase (*OTC*), urea transport (*Urt*), arginase (*Arg*), citrate synthase (*Glta*), photosystem I (*Psa*), photosystem II (*Psb*), cytochrome c oxidase (*COX*), cytochrome b (*PetB*), and cytochrome c (*PetC*).

Evaluating these physiological trends further, our findings appear consistent with those previously reported for the closely-related chlorophyte, *Ostreococcus* (Lelandis et al., 2016), with *Micromonas* appearing to contain far fewer transcribed genes involved in nitrogen and iron metabolism than are commonly reported in diatoms and other phytoplankton taxa. Transcripts for the nitrogen assimilation genes nitrate/nitrite transporters (*NRT*), nitrate reductase (*NR*), nitrite reductase (*NiR*), ammonium transporters (*AMT*), and urea transporters (*Urt*) were identified in our *Micromonas* isolate (Fig. 4B, 4C). Most genes associated with NO_3_^−^ uptake (*NRT*, *NiR*) increased expression (Log2Fold changes > 0) in the Fe+ treatment relative to Fe− treatment (Fig. 4B). Urea transporters were also overrepresented (Log2Fold change = 4.21; adjusted p-value < 0.01) under Fe+ conditions. Conversely, ammonium transporters showed a slight negative Log2Fold change (−0.47), suggesting increased expression in the Fe− treatment (Fig. 4B, 4C). Overall, genes associated with the expression of photosystems I and II (*Psa*, *Psb*) exhibited a stronger Log2Fold change (−2.93) in Fe− conditions, in contrast to the common assumption that these photosynthetic genes should be more highly expressed under iron-replete conditions (Fig. 4B).

Our data suggests that *Micromonas* may lack genes commonly involved in iron regulation and homeostasis in diatoms residing in HNLC regions, including proton-pumping rhodopsins and ferritin (Marchetti et al., 2009; Andrew et al., 2023). However, we did identify the gene for iron starvation induced protein 3 (*ISIP3*) which was strongly expressed (Log2Fold change = −4.62; adjusted p-value <0.01) under Fe− conditions (Fig. 4B). The percent of overall contigs without a functional annotation was high (87.9%), suggesting additional genes involved in both nitrogen and iron metabolism may be present but are not currently identified in the KEGG database.

## 4. Discussion

### 4.1 Cellular and metabolic characterization

While less is known about the ecology and physiology of *Micromonas* relative to other picoeukaryotic chlorophytes such as *Ostreoccocus*, our findings reported here are consistent with and build upon some of what has been reported on *Micromonas* (Cochlan and Harrison, 1991a; 1991b). Our estimates of cell size are similar to the previously reported values of <2 µm (McKie-Krisberg and Sanders, 2014). Additionally, *Ostreoccocus* has been reported to exhibit fast growth rates under nitrogen replete conditions (µ_max_ of 1.11 d^−1^; Cochlan and Harrison, 1991a), consistent with our measured growth rates for our *Micromonas* sp. isolate in iron-replete conditions. However, a substantial lag phase has not been previously reported. Additionally, our inability to achieve steady state growth in our isolate using NH_4_^+^ media is noteworthy. In fact, following nitrogen starvation, Cochlan and Harrison (1991a; 1991b) reported a preference for NH_4_^+^ and urea uptake relative to NO_3_^−^ uptake. While we were unable to identify what was inhibiting growth, it is likely that our *Micromonas* sp. requires specific conditions for NH_4_^+^ utilization that were not met within our culture conditions. Our findings suggest that our strain of *Micromonas* may exhibit a preference for NO_3_^−^ over other forms of nitrogen (such as NH_4_^+^), but further research is warranted.

Our study quantified changes in cellular Chl *a* concentration and C and N quotas in iron-replete and deplete *Micromonas sp.* cells. Understanding these iron limitation-driven differences is critical as changes in growth rates and elemental quotas of *Micromonas* may have ecosystem and biogeochemical implications in HNLC regions. Consistent with current paradigms of diatoms, iron-replete *Micromonas* sp. were larger, had higher growth rates, higher cellular Chl *a* content, and appreciably higher C and N quotas relative to their iron-limited counterparts. Although the Fe− limited cells were smaller with lower growth rates and cellular quotas, they were able to reach higher cell densities under similar culturing conditions. These findings suggest *Micromonas* can achieve numerically dominance in iron-limited environments through reducing their cellular elemental quotas.

### 4.2 Iron limitation strategies

Our findings are in line with previous work (Martin and Fitzwater, 1988; Boyd et al., 1998; Marchetti et al., 2006) suggesting low Fe concentrations limit nitrate uptake as is supported by a low relative growth rate (µ/µ_max_). This leads to substantially curtailed growth rates, and trends in gene expression where genes involved in nitrate metabolism (e.g., *NRT*, *NiR*) are more highly expressed under iron-replete conditions. The enhanced expression of genes related to nitrogen metabolism under iron replete conditions as well as our Fe+ modeled rates consistent with those small celled new production rates measured during EXPORTS suggests that picoeukaryotes such as *Micromonas* may be an important contributor to new production in the iron-limited waters of the Northeast Pacific. Our data also demonstrates clear expression patterns in genes involved in NO_3_^−^ metabolism coinciding with their ability to grow effectively using NO_3_^−^.

*Micromonas* also appears capable of urea metabolism by expressing urea transporters in addition to *NiR* and *NRT*. A similar finding was reported by Piganeau et al. (2011) who exhibited selective pressure for nitrate and urea uptake, but not ammonium uptake. This finding is consistent with our inability to establish steady-state growth when cells were grown in NH_4_^+^ based media. However, this is surprising given the need to reduce NO_3_^−^ to NH_4_^+^ for utilization in protein synthesis and the lower energetic cost of NH_4_^+^ utilization relative to NO_3_^−^ utilization (Raven, 1988). Cultures were able to grow sporadically on NH_4_^+^, but steady-state growth was not achieved. This, combined with the presence of *AMT* in the transcriptome, supports previous studies (Cochlan and Harrison, 1991a; 1991b) and suggests *Micromonas* have the physiological capabilities for NH_4_^+^ utilization but were perhaps inhibited by some other factor, such as a vitamin or a micronutrient.

The lack of information on iron regulation is consistent with findings from previous studies of the closely related *Ostreococcus* (Sutak et al., 2012; Lelandais et al., 2016). The majority of what is known about phytoplankton iron metabolism comes from studies of diatoms (Peers and Price, 2006; Marchetti et al., 2012). It is therefore not surprising that many of the iron transport and homeostasis genes identified in diatoms were not readily identified in *Micromonas* given the substantial physiological differences between the two phytoplankton groups. The only iron-related gene identified and present in the *Micromonas* transcriptome is ISIP3, an iron-starvation induced protein with ferritin-like domains (Behnke and LaRoche, 2020). However, as is noted by Lelandais et al. (2016) for *Ostreococcus*, the high percentage of unannotated genes in our strain suggests that chlorophytes likely have novel methods for dealing with iron limitation which have not yet been well characterized. Sutak et al. (2012) also found both *Ostreococcus* and *Micromonas* were able to take up ferric and ferrous iron without prior iron reduction but unable to store it. The authors hypothesize ferritin may be present in *Ostreococcus* but not in *Micromonas* (Sutak et al., 2012). Our study represents the first analysis of *Micromonas* isolated from an HNLC region. The cosmopolitan nature and wide thermal niche of *Micromonas* suggests it can adapt to new or changing environments and proliferate quickly (Vannier et al., 2016; Demory et al., 2018). This may suggest the isolate studied here exhibits physiological characteristics and adaptations that may be quite different from what is currently characterized in the scientific literature using isolates from other regions.

An additional, noteworthy finding is that our data appears to suggest that *Micromonas* exhibit a form of nitrogen metabolism conservation even under Fe replete conditions. The gene expression analysis demonstrates most protein-encoding genes related to nitrogen metabolism (*NRT*, *NR*, *AMT*, *nirA*) have some of the highest constitutive transcript abundances of any pathway, exhibiting some significant directionality of gene expression under variable Fe conditions, but only to a minimal extent (Fig. 4B). Some studies have suggested diatoms exhibit substantial changes in expression of genes involved in nitrogen metabolism, particularly in the natural environment (Marchetti et al., 2012), whereas others have likewise shown a lack of substantial change in gene expression (but not protein synthesis) in diatoms grown in culture (Cohen et al., 2018). This finding supports previous studies which suggest picoeukaryotes, and other small phytoplankton, provide a “baseline” rate of primary production under limiting conditions with a minimal physiological response to dramatic shifts in environmental conditions (Meyer et al., 2022). It is possible that *Micromonas* expresses additional genomic adaptation strategies for iron limitation that were not identified from our transcriptome analysis. Additional research is needed to fully elucidate the strategies *Micromonas* uses to subsist in iron-limited environments.

### 4.3 Oceanographic context

Our study provides valuable cellular characteristics of an abundant, picoeukaryote in a climatologically important oceanographic context. Meyer et al. (2022) found that small cells (<5 µm) accounted for the majority of biomass (68% of total biomass) and primary production (89% of total primary production), including new production (72% of total new production), at Station P during the EXPORTS observation period. Furthermore, *Micromonas* appears to be highly represented within the small-celled phytoplankton community, with 18S rDNA data indicating that *Micromonas* was within the top 20% of the most abundant phytoplankton genera and one of the top five most abundant picoeukaryotes (Jones and Rynearson, 2022).

Our estimated rates of new production by picoeukaryotes such as *Micromonas*, taken as a representative species, under Fe− and Fe+ conditions were compared to the average small-celled new production rates of 33.3 ± 14.4 nmol N L^−1^ d^−1^ from the NP EXPORTS campaign (Fig. 3). While higher rates of NO_3_^−^ uptake may occur within a culture relative to a mixed, natural phytoplankton community, our analysis supports the idea that *Micromonas* contributes substantially to total new production at Station P, especially when iron concentrations can support higher growth rates and cellular nitrogen quotas. Inherently, other phytoplankton groups, both within the picoplankton and nanoplankton size ranges, present at St. P during the EXPORTS observation period also contribute appreciably to new production. Our culture-based analysis is not meant to suggest *Micromonas* accounted for all of this new production, but rather, that this chlorophyte should at least be considered as a significant contributor to new production (and by inference carbon export) as estimated from in situ picoeukaryote cell densities along with our measured cellular nitrogen quotas and growth rates as a function of iron status.

With projected decreases in euphotic zone nutrient concentrations in many parts of the oceans as a result of climate change (Kwiatkowski et al., 2020), induced changes in water mass stratification and overall circulation, models of future projections predict an overall shift towards dominance by smaller phytoplankton groups such as chlorophytes (Henson et al., 2021). Abundant, small-cell phytoplankton with streamlined genomes and enhanced capabilities to adapt to changing environmental conditions such as *Micromonas* hold distinct advantages in future climate conditions. Further studies on *Micromonas* grazing rates, sinking speeds, etc. are needed to fully understand its ecology moving forward and how this will impact net carbon export rates and trophic level interactions.

## Conclusion

In low iron environments, *Micromonas sp.* exhibits physiological strategies, including reduced growth rates, small cell size, and metabolic restructuring, to combat Fe limitation and maintain high rates of new production and substantial abundances. This study furthers our understanding of this poorly characterized picoeukaryote and verifies a potential role for chlorophytes in performing new production in iron-limited regions. Here, we add to the body of knowledge reaffirming the need to shift our thinking on key phytoplankton groups that have the potential to contribute to carbon export and pay additional attention to the role picoeukaryotes play in ecosystem dynamics both in current and future ocean climate scenarios.

## Acknowledgements

We thank the EXPORTS Chief Scientists, D. Steinberg and J. Graff, and the Captain and crew of the *R/V* Roger Revelle for their roles in the initial sample collection, R. Paerl for use of his flow cytometer, and J. Graff for NP EXPORTS flow cytometry data. We also thank R. Flynn for her helpful comments. This work was supported by NASA Grant 80NSSC17K0552 and NSF Grant OCE2219973 to AM.

## Competing Interests

The authors declare no competing interests.

## Data Availability Statement

All data presented here are publicly available at the National Center for Biotechnology Information (Accession number PRJNA1072555) or the Biological and Chemical Oceanography Data Management Office (project ID: 914269).

## Notes

https://www.bco-dmo.org/project/914269

https://www.ncbi.nlm.nih.gov/sra/SRX23571111[accn]

### Competing Interest Statement

The authors have declared no competing interest.

### Summary of Updates

This version of the manuscript has been revised to reflect updated 18S identification from the National Center for Biotechnology Information.

